# Bacteriophage FNU1 negates *Fusobacterium nucleatum* induced cell growth and chemotherapy resistance in gastrointestinal cancer cells

**DOI:** 10.1101/2025.07.10.664266

**Authors:** Mwila Kabwe, Birhanu Ayelign, Shoukat Afshar-Sterle, Michael Buchert, Joseph Tucci

## Abstract

*Fusobacterium nucleatum* is an oncobacterium capable of promoting the growth and chemotherapy resistance of colonised tumours. Although *F. nucleatum* is usually susceptible to a range of antibiotics, these have been associated with worse outcomes when administered with anti-neoplastic chemotherapy. Bacteriophages are viewed as natural alternatives to antibiotics that provide bacterial-specific targeting. In this study, we demonstrated that gastric tumour tissues and cells expressed Gal-GalNac molecules that facilitate binding of the fap2 receptor on *F. nucleatum*, leading to increased cancer cell proliferation and migratory potential. We then employed an *F. nucleatum* specific bacteriophage, FNU1, to limit the effects of this oncobacteria in colon cancer and gastric cancer cell models. We demonstrated that FNU1 was able to negate the *F. nucleatum*-dependent growth stimulatory effects, migratory ability, induction of reactive oxygen species, and autophagy in these cell lines. *F. nucleatum* also inhibited apoptosis in co-cultures with colon and gastric cancer cells, and FNU1 acted synergistically with the chemotherapy agents 5-fluorouracil and oxaliplatin to induce apoptosis in these models. Treatments with bacteriophage FNU1, therefore, have the potential to augment existing cancer therapy, and further testing in animal models is warranted.

## Introduction

Cancer is one of the leading causes of death worldwide [1]. The gastrointestinal tract (GIT), which includes the oral cavity, pharynx, and other regions of the digestive tract, includes tissues and organs with the highest incidence of cancers, some of which are the leading cause of cancer-related deaths globally [2,3]. The GIT is a unique system eliciting innate and adaptive immune functions to keep out pathogens, as well as facilitating nutrient absorption [4]. The symbiosis between the GIT and its microbiota is important in immune tolerance, immune modulation, and maintenance of metabolic homeostasis [5]. The microbiome of the GIT is composed of bacteria and viruses, and dysbiosis in these tissues has been associated with changes in the metabolome, immune dysregulation, and cancer [6,7]. Both, bacteria and eukaryotic viruses have pathogenic mechanisms that have been linked directly to carcinogenesis [6,8] whilst bacterial viruses (bacteriophages) regulate bacterial populations leading to a dynamic [7] and potentially manipulable microbiome. This characteristic of bacteriophages makes them an attractive resource for the potential manipulation of the microbiome in a dysbiotic state.

Several genomic sequencing data have shown a select few microbes are increased in tumour tissues [9]. Among several cancers, the bacteria *Fusobacterium* is the most frequently increased and the most widely studied, especially in the context of colon cancers. *Fusobacterium nucleatum* is an heterogenous oral pathobiont [10] known to promote tumourigenesis by dampening the host’s anti-tumour immunity, promoting the release of pro-inflammatory cytokines and by directly stimulating the proliferation of tumour cells [9]. The interactions between *F. nucleatum* and colon cancer cells are complex. Studies have highlighted that through the TLR4 receptor, *F. nucleatum* increases proliferation of colon cancer cells [11], inhibits apoptosis by upregulation of BIRC3 [Baculoviral IAP (Inhibitor of Apoptosis Protein) Repeat-Containing 3] [12], and modulates autophagy to promote chemotherapy resistance [13]. Usually, in epithelial cells, DNA damage-induced apoptosis is promoted by the caspase-3 enzyme through proteolysis of the mismatch repair protein MLH1 (MutL homolog 1) [14]. However, in the presence of *F. nucleatum*, reactive oxygen species (ROS) are produced [15] that cause epigenetic silencing of MLH1, leading to aberrant DNA methylation and microsatellite instability [16,17] to inhibit apoptosis [14].

The use of antibiotics to control *F. nucleatum* in cancer has been examined [18]. However, antibiotic treatment before chemotherapy has been associated with worse outcomes in mouse models [19] and in humans is associated with disease progression and shortened survival [20,21]. Further, a history of antibiotic use as people age has been associated with an increased risk of development of GIT cancers [22-25] and the prevalence of *F. nucleatum* in tumourigenic tissues has been shown to be independent of antibiotic use [26]. Recent JAVELIN and KEYNOTE-061 phase III international clinical trials have shown that GIT cancers do not respond to immunotherapy [25,27,28] and evidence has suggested this may be a result of the human microbiota’s impact on gut inflammation and immune dysregulation [29]. While the potential for using bacteriophages against key oncobacteria in augmenting cancer therapies has been reviewed [9], there has not been clear evidence to support how bacteriophages can modulate the *Fusobacterium* infection of gastric and colon cancer cells to elicit their anti-cancer benefits.

## Materials and Methods

### Ethics approval

All study protocols have been approved by the La Trobe University Ethics Committee (S17-112) and all methods performed in accordance with the La Trobe University Ethics, Biosafety and Integrity guidelines, and regulations.

### Bacteria culture conditions and Bacteriophage purification

The fully characterised *F. nucleatum* strain (ATCC10953) [30] and bacteriophage FNU1 (Accession number: MK554696) were used to elucidate the interactions between bacteria, bacteriophage, and cancer cells. The bacteria were cultured in Brain Heart Infusion (BHI; Oxoid^™^, Australia) media supplemented with 0.5% cysteine (Sigma-Aldrich^®^, Australia) and 0.5% haemin (Sigma-Aldrich^®^, Australia). The bacterial cultures were incubated at 37^°^Cunder anaerobic conditions using AnaeroGen pack (Oxoid^™^, Australia) [31].

The broth macrodilution method [32] was used to determine the minimum inhibitory concentration of ampicillin against *F. nucleatum*. Ampicillin was serially diluted in BHI broth containing 5×10^5^ CFU/mL. The dilutions were then plated onto BHI agar plates and incubated for 48 h anaerobically at 37 ^°^C before performing colony counts.

Bacteriophage FNU1 was clonally expanded from one plaque and purified by 0.2 µm filtration. The concentrated filtered stock was precipitated in sodium chloride (NaCl) and polyethylene-glycol (PEG 8000) (Sigma-Aldrich^®^, Australia) and treated with 2 % v/v Triton^™^ X-100 (Sigma-Aldrich^®^, Australia) [33] before washing three times in phosphate-buffered saline (PBS; pH 7.4) and resuspended in 10 % FBS supplemented RPMI-1640 medium containing L-glutamine and sodium bicarbonate (Sigma-Aldrich^®^, Australia) [34]. Bacteriophages were added to the *F. nucleatum*/cancer cell co-culture at a MOI of 0.1 (with reference to *F. nucleatum* cells).

### Human cell line maintenance

Colon HCT116 cells (ATCC, USA) and gastric MKN1 (ATCC, USA) cancer cell lines purchased from the American Type Culture Collection were maintained in 10% (v/v) fetal bovine serum (FBS; Sigma-Aldrich^®^, Australia) supplemented RPMI-1640 medium containing L-glutamine and sodium bicarbonate (Sigma-Aldrich^®^, Australia). Cells were passaged by detachment with 1× trypsin-EDTA solution (0.05% trypsin, 0.02% EDTA, in Hanks′ Balanced Salt Solution (Sigma-Aldrich^®^) and maintained in humidified 5% CO_2_ at 37 ^°^C. Colon and gastric cancer cell lines were grown up to 90% confluence before harvesting with 1× trypsin and resuspending in fresh 10% (v/v) FBS supplemented RPMI-1640 medium containing L-glutamine and sodium bicarbonate (Sigma-Aldrich^®^, Australia) to a concentration of 1×10^4^ cells per mL.

### Bacterial/bacteriophage and cancer cells co-cultures

Colon and gastric cancer cell lines were grown up to 90 % confluence before harvesting with 1× trypsin and resuspending in fresh 10 % FBS supplemented RPMI-1640 medium containing L-glutamine and sodium bicarbonate (Sigma-Aldrich^®^, Australia) to a concentration of 1×10^4^ cells per mL.

Exponentially growing bacteria in BHI broth were harvested to a concentration of 1×10^8^ colony forming units (CFU) per mL. One mL containing 1×10^8^ cells of *F. nucleatum* were washed 3 times by centrifugation at 12, 000 × ℊ and resuspended in RPMI-1640 medium containing L-glutamine and sodium bicarbonate (Sigma-Aldrich^®^, Australia). *F. nucleatum* in RPMI-1640 was then added to the 1×10^4^ cells per mL of cancer cells at a MOI of 10.

*F. nucleatum*/cancer cell/FNU1 co-cultures were incubated in humidified 5 % CO_2_ at 37 ^°^C for 24 h. The *F. nucleatum* infection was confirmed by passaging the co-culture and then growing it on BHI agar to observe colonies as well as through PCR of 16s rRNA primers (U27F: 5’ - AGAGTTTGATCMTGGCTCAG – 3’ and U492R: 5’ - AAGGAGGTGWTCCARCC - 3’) under thermocycling conditions: 95 ^°^C for 3 min; 32 cycles of 95 ^°^C for 30 s, 60 ^°^C for 30 s, and 72 ^°^C for 90 s; then a final extension at 72 ^°^C for 10 min [31] and *F. nucleatum* specific primers (5’ – AGTTGTCCTATACCAGCTCCAAC - 3’ and 5’ – GCAACATTCTTTGCAGCACGTACTGC – 3’) under thermocycling conditions: 95 ^°^C for 10 min; 35 cycles of 95 ^°^C for 30 s, 60 ^°^C for 30 s, and 72 ^°^C for 45 s; then a final extension at 72 ^°^C for 10 min.

### Immunofluorescence microscopy

Human gastric MKN1 cells and colorectal HCT116 tumour cells were seeded in 24 well plates and after overnight culture, cells were washed three times with RPMI 1640 cell growth medium and fixed with 4 % paraformaldehyde. Cells were blocked with 1% of BSA for 30 min at room temperature. Fluorescein isothiocyanate conjugated lectin (FITC-lectin, specific for the sugar molecule D-galactose-β (1–3)-N-acetyl-D-galactosamine (Gal-GalNAc), ThermoFisher Scientific, Australia) was then incubated (50 μg/mL in PBS) overnight at 4 ^°^C, followed by staining with Hoechst 33342 (1:1000 of PBS) for 30 min at room temperature. Finally, the slides were mounted with Fluoromount-G^™^ Mounting Medium (Invitrogen, ThermoFisher Scientific, Australia). Images were visualized using Zeiss LSM 980 Axio Observer 7 inverted confocal microscope.

### Immunohistochemistry

Mouse gastric tumour tissue was provided by Dr Moritz Eissmann, Cancer and Inflammatory Laboratory, Olivia Newton Cancer Research Institute, Heidelberg, Australia. The slides were deparaffinized using xylene for 15 min twice, 100 % alcohol twice and 70 % alcohol for 5 min, then treated with antigen retrieval solution (10mM of sodium citrate pH 6) in a microwave for 20 min and then left for 40 min at room temperature, followed by a final wash for 5 min in PBS to remove any impurities. Non-specific binding was minimised by blocking the slides with a solution of PBS supplemented with 10 % FBS, 10 % BSA, and 5 % Triton-X for one hour at room temperature. Slides were then incubated overnight at 4 ^°^C in 50 μg/mL in PBS of FITC-conjugated lectin followed by incubation with Hoechst 33342 (1:1000 of PBS) for 30 min at room temperature. Finally, the slides were mounted with Fluoromount-G^™^ Mounting Medium (Invitrogen, ThermoFisher Scientific, Australia) and visualised using an immunofluorescence microscope.

### Flow cytometry analysis

The expression level of Gal-GalNac in *<u>K</u>ras*^*G12D*^, *<u>P</u>i3kCA*^*H1047R*^ and *<u>T</u>p53*^*R172H*^ (KPT) mutant murine gastric cancer organoids and human gastrointestinal cancer cells was performed using flow cytometry. To obtain single cells, murine cancer organoids were dissociated from the Matrigel using 1× trypsin with EDTA (0.05 % trypsin, 0.02 % EDTA). Briefly, cells were enzymatically detached with 1× trypsin with EDTA and cells were washed three times with RPMI medium. Cells were blocked with 1 % of BSA for 30 min at room temperature. Single cells were stained with FITC-conjugated lectin at a concentration of 50 µg/mL in PBS for 30 min at room temperature and then washed three times in PBS. The cells were stained with Sytox blue before being analysed by flow cytometry. The data was analysed using FlowJo 10.10 software.

### Laser scanning confocal microscopy (LSCM) and *F. nucleatum* invasion

Exponentially growing *F. nucleatum* was adjusted to 1×10^8^ CFU/mL in PBS (pH 7.4) and stained with the CellTrace^™^ CFSE cell proliferation reagent according to manufacturer instructions (Invitrogen^™^, Australia). Briefly, 2 µl/mL were added to the *F. nucleatum* suspension and incubated for 30 min. CellTrace^™^ CFSE stained *F. nucleatum* were resuspended in fresh 10 % (v/v) FBS supplemented RPMI-1640 medium containing L-glutamine and sodium bicarbonate (Sigma-Aldrich^®^, Australia) and added to 1×10^4^ cells per mL of cancer cells at MOI of 10. The co-culture was then transferred to 6-well plates with coverslips and incubated in humidified 5 % CO_2_ at 37 ^°^C for 24 h. Unattached bacteria were washed off in fresh RPMI-1640 media twice before adding fresh media containing bacteriophage FNU1 (MOI = 0.1 to initial starting concentration of *F. nucleatum*) or ampicillin (1mg/mL; Sigma-Aldrich^®^, Australia) and incubating a further 2 h in humidified 5 % CO_2_ at 37 ^°^C. The RPMI-1640 media was then removed, and excess media washed off three times in PBS before staining cell nucleus with Hoechst 33342 (2 µl/mL; Sigma-Aldrich^®^, Australia) and cytoplasm with CellBrite^®^ Red cytoplasmic membrane dye (2 µl/mL; Biotium, Australia). The cells were then incubated for 30 min before washing off in fresh RPMI-1640 media and incubating a further 5 min at 37 ^°^C. The RPMI medium was then washed off in PBS and the cover slips mounted on microscope slides with PBS, sealed with nail polish and examined immediately under the Olympus Fluoview Fv10i confocal laser scanning microscope (Olympus Life Science, Australia). The three-channel excitation and emission (Ex/Em) wavelengths were 492/517 nm (CellTrace^™^ CFSE), 644/665 nm (CellBrite^®^ Red) and 361/497 nm (Hoechst 33342).

### Cell Proliferation Assay

Cell proliferation was quantified using the Sulforhodamine B (SRB) assay [35]. Co-cultures treated with 5-fluorouracil (5-FU, 20 µM) and oxaliplatin (20 µM) were incubated in humidified 5 % CO_2_ at 37 ^°^C for 24 h before fixation in 10 % (w/v) trichloroacetic acid (ThermoFisher Scientific, Australia) at 4 ^°^C for 30 min. The trichloroacetic acid was gently pipetted off and fixed cells washed in milli-Q water (Merck Milli-Q water system, Australia) five times and air dried at room temperature. The cells were then stained with 1 % (w/v) SRB (Sigma-Aldrich^®^, Australia) in glacial acetic acid (ThermoFisher Scientific, Australia) for 15 min. The SRB was then pipetted off and excess stain washed off in 1 % (v/v) glacial acetic acid and air-dried. SRB that remained bound to the cells was solubilised in 10 mM unbuffered TRIS base and its absorbance quantified using the FlexStation 3 plate reader (Molecular Devices, United States).

### Wound healing migration assay

To determine the effect of *F. nucleatum* infection and FNU1 bacteriophage treatment on migration of MKN1 gastric cancer cells and HCT116 colon cancer cells, we performed wound healing or scratch assays under different conditions: (i) uninfected cells, (ii) uninfected cells treated with FNU1, (iii) *F. nucleatum* infected GI cancer cells, and (iv) *F. nucleatum* infected cells treated with FNU1. Migration was quantified by measuring the percentage of wound closure at 24-, 48- and 72-h post-wound scratch. Wound healing assays were performed according to the protocol used by Ou and colleagues [36]. In brief, 1 × 10^5^ MKN1 cells and HCT116 cells were seeded in 6-well plates and incubated at 37 ^°^ C and grown to 100 % confluence. The cells of each well were then infected with *F. nucleatum* at 1:30 MOI, or equal volume of PBS as a control, and incubated for 2 h. The cell monolayers were then scratched with 20 µl pipette tip in a straight line to create a “wound” followed by two washes with PBS and incubation for 24 h by adding serum-free medium and 1:10 MOI of FNU bacteriophage. Images of the same scratched area were taken using a ZeissAxio observer widefield microscope at 0, 24, 48 and 72 h. The scratched area was measured by ImageJ software and used to calculate wound closure percentage and rate.

### Intracellular reactive oxygen species (ROS) quantification

Intracellular ROS production was quantified using the DCFDA cellular ROS detection kit (Abcam^®^, Australia) according to the manufacturer’s instructions. Briefly, cancer cells were grown to 90 % confluence and harvested by trypsinisation before washing and resuspending in fresh phenol free RPMI-1640 media containing 20 µM DCFDA solution. The cell suspension was incubated in the dark at 37 ^°^C for 30 min before resuspending in fresh phenol free RPMI-1640 media containing 20 µM of the drugs 5 – fluorouracil or oxaliplatin at a concentration of 1×10^4^ cells/mL. The cells were then seeded into sterile, dark, clear bottom 96-well microplates to which *F. nucleatum* at MOI of 10 and bacteriophage FNU1 at MOI of 0.1 was added. Whilst incubating in humidified 5 % CO_2_ at 37 ^°^C, fluorescence was read every 2 h for 6 h at Ex/Em of 485/535 nm using the FlexStation 3 plate reader (Molecular Devices, United States). Tert-butyl hydrogen peroxide solution (20 µM) was used as the assay positive control.

### Autophagy quantification

Autophagy was assessed using the autophagy detection kit (Abcam^®^, Australia) according to the manufacturer’s instructions. Co-cultures were grown for 24 h in the same manner as described in the cell proliferation assay, except in this case, using phenol free cell growth media in sterile, dark, clear bottom 96-well microplates. Rapamycin (500 nM) was added at the start of the co-culture as a positive control; 3 h prior to assaying for autophagy, media for cancer cells in negative control wells was replaced with Earle’s balanced salt solution (ThermoFisher Scientific, Australia) with 20 µM chloroquine and re-incubated in humidified 5 % CO_2_ at 37 ^°^C. All media was then removed from wells, and cells washed twice in 1× assay buffer before staining with the kit’s green detection reagent (2µl/mL) and incubating at 37 ^°^C for 30 min. The stain was then removed. Cells were washed twice with 1× assay buffer to remove excess stain before adding 100 µL of 1× assay buffer to each well and quantifying fluorescence at Ex/Em of 465/534 using the FlexStation 3 plate reader (Molecular Devices, United States).

### Apoptosis assay

Cellular apoptosis was evaluated using the Beckman Coulter CytoFLEX Research Flow Cytometer (Beckman Coulter^®^ Life Sciences, Australia) after staining with Annexin V and Propidium iodide (PI), TACS^®^ Annexin V-FITC Apoptosis Detection kit (Trevigen^®^, United States). Co-cultures of cancer cells with bacteria and/or bacteriophages were incubated for 24 h before replacing media with fresh media containing 5-fluorouracil or oxaliplatin at concentration of 100 µM. These were then incubated for a further 24 h in humidified 5 % CO_2_ at 37 ^°^C. The cells were then harvested by trypsinisation and washed twice in PBS before staining with PI (2µl/mL) and Annexin V (2µl/mL) in 1× Annexin Binding buffer (ThermoFisher Scientific, Australia). This was incubated in the dark at room temperature for 30 min before capturing 50 000 events by flow cytometry. Data were analysed in the CytExpert software version 2.3.1.22 (Beckman Coulter, Inc., Australia). The proportion of apoptotic and necrotic cells were determined by adding the upper right quadrant (late apoptotic, secondary necrosis), lower left quadrant (early apoptotic – phosphatidylserine exposed, membrane intact) and upper left quadrant (necrotic - membrane compromised, not apoptotic).

### Statistical analysis

All data were imported into and analysed in the IBM^®^ SPSS^®^ software platform version 27 (SPSS, Inc., United States). Normality of quantitative data were determined by the Shapiro-Wilk test with the paired *t-*test being used to compare means between two groups with normally distributed data while the non-parametric counterpart, Wilcoxon signed-rank test was used for data that significantly deviated from the normal distribution. For two or more groups of data, we compared their means using Analysis of Variance. For all tests, the *p* value of less than 0.05 was considered statistically significant.

## Results

### Widespread expression of Gal-GalNac in human GI cancer cells mediates *F. nucleatum* infection

It has been shown that the presence and overexpression of the sugar molecule Gal-GalNac in gastrointestinal cancer cells facilitates the binding and cellular invasion of *F. nucleatum* through its interaction with its protein, *Fusobacterium* adhesin protein 2 (Fap2), and allows enrichment of *F. nucleatum* at tumour sites [37].

In gastric tumour tissues (corpus and antrum), we found a significantly higher expression of Gal-GalNAc compared to the wildtype tissues (*p<0*.*01*) (Figure 1A and B) in the *gp130*^*F/F*^ mouse model of inflammatory cytokine-driven intestinal metaplasia and early gastric cancer that affects both antrum and corpus regions of the murine stomach [38]. In this model, there was a higher expression of Gal-GalNAc in the antrum compared to the corpus tissues, *p < 0*.*001*. We also found that Gal-GalNAc expression was seen in more than 60 % of *<u>K</u>ras*^*G12D*^, *<u>P</u>i3kCA*^*H1047R*^ and *<u>T</u>p53*^*R172H*^ (KPT) mutant murine cancer organoids [39], irrespective of whether the cancer organoids were maintained ex vivo in 3D static cultures or grown as orthotopic tumours in vivo (Figure 1C and D).

**Figure 1:**
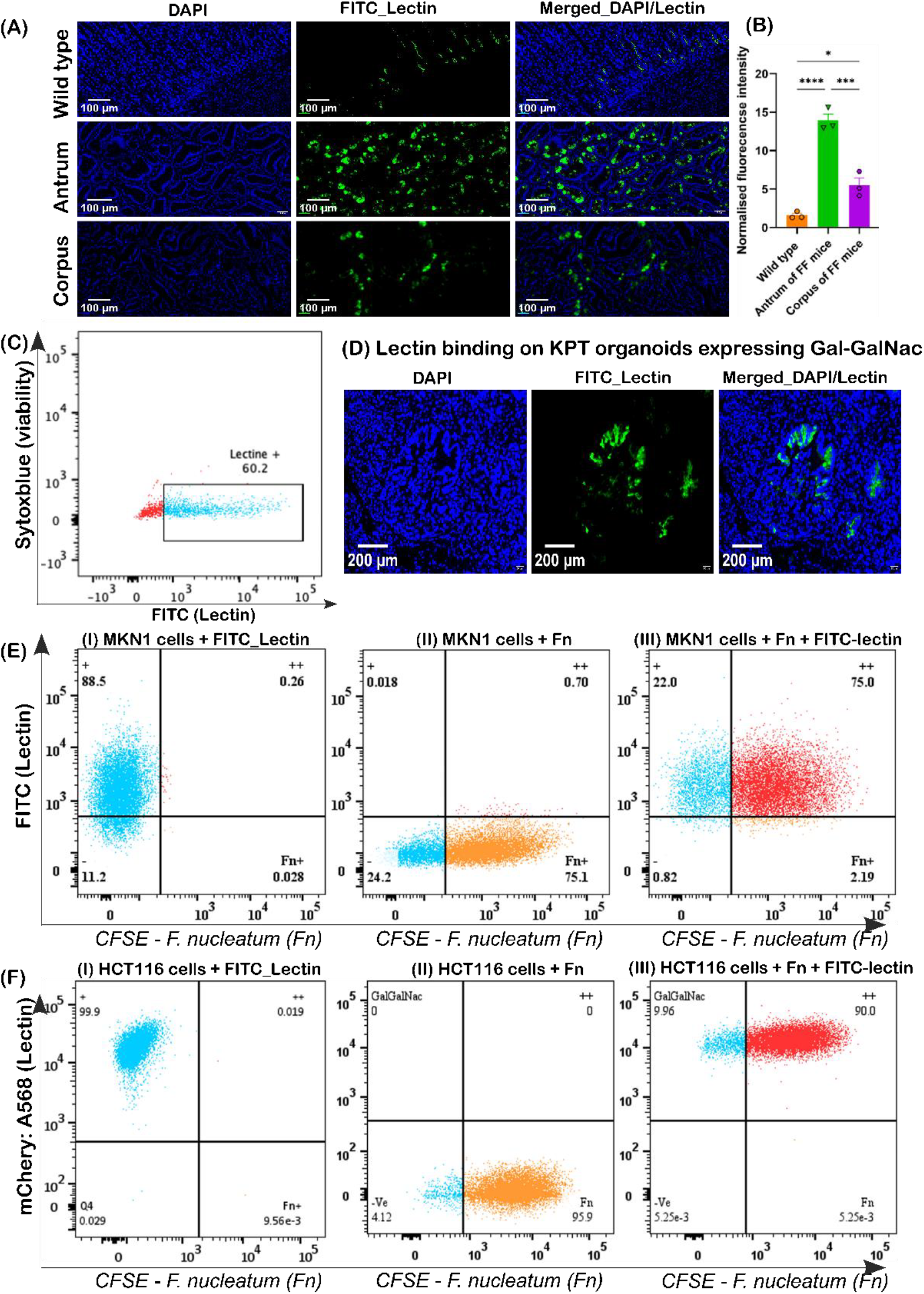
Co-localisation of Gal-GalNac expression and F. nucleatum infected gastrointestinal tumour cells. A) Visualisation of Lectin bound to gastric corpus and antrum tumour tissues in gp130^F/F^ (FF) mice. B) Significantly higher expression of Gal-GalNAc in tumour antrum compared to corpus, p value <0.01, and both greater than wild type tissue, p-value <0.001. C) Gal-GalNAc expression in KPT murine gastric cancer organoids and in D) orthotopic KPT tumours visualized using confocal microscope. DAPI fluorescent dye stains chromosomes blue. E) Flow cytometry analysis of Gal-GalNac expression in MKN1 cells infected with cellTrace™ far-red labelled F. nucleatum, multiplicity of infection 1:10. F) Flow cytometry analysis of Gal-GalNac expression in HCT116 cells infected with cellTrace™ CSFE labelled F. nucleatum, multiplicity of infection 1:30 (Fn: F. nucleatum).

Similarly, we evaluated the level of Gal-GalNac expression in two human GI cancer lines, the gastric adenocarcinoma cell line MKN1 and the colon carcinoma cell line HCT116. Flow cytometry of cells stained with a Gal-GalNac-specific, FITC-labelled lectin indicated that more than 90 % of cells displayed Gal-GalNac sugar moieties on their cell surface [Figure 1E(I) and 1F(I)]. 24 h after infection with fluorescently labelled *F. nucleatum*, close to 90 % of *F. nucleatum* positive cells [Figure 1E(II) and 1F(II)] also stained positive for lectin [Figure 1E(III) and 1F(III)] suggesting that *F. nucleatum* interaction with MKN1 and HCT116 cells is Gal-GalNac dependent.

### FNU1 bacteriophage eliminates intracellular *F. nucleatum*

Using laser scanning confocal microscopy, we were able to visualise GI cancer cells (marked by blue staining of cytoplasm and cyan staining of nucleus) (Figure 2); when co-cultured with *F. nucleatum* (magenta staining), we observed that *F. nucleatum* cells existed extracellularly and intracellularly [Figure 2 A(ii) and B (ii)]. When the co-culture was treated with the bacteriophage FNU1, both intracellular and extracellular *F. nucleatum* were eliminated [Figure 2 A(iii) and B (iii)]. Initially, minimum inhibitory concentration of the antibiotic ampicillin against *F. nucleatum* using BHI broth dilution was determined at 2 µg/mL. When the co-culture was treated with 1 mg/mL ampicillin the antibiotic was able to eliminate extracellular bacteria but not the intracellular *F. nucleatum* [Figure 2 A(iv) and B (iv)]. After passaging we showed that *F. nucleatum* remained viable in the co-culture (Figure 2C, left side) and following inoculating on BHI agar, observed a significant increase (*p* < 0.001) in the colony counts of *F. nucleatum* between the first and second passage of the co-cultures. There were no colonies observed after the first and second passages in bacteriophage treated co-cultures of GI cancer cells (Figure 2C, right side).

**Figure 2:**
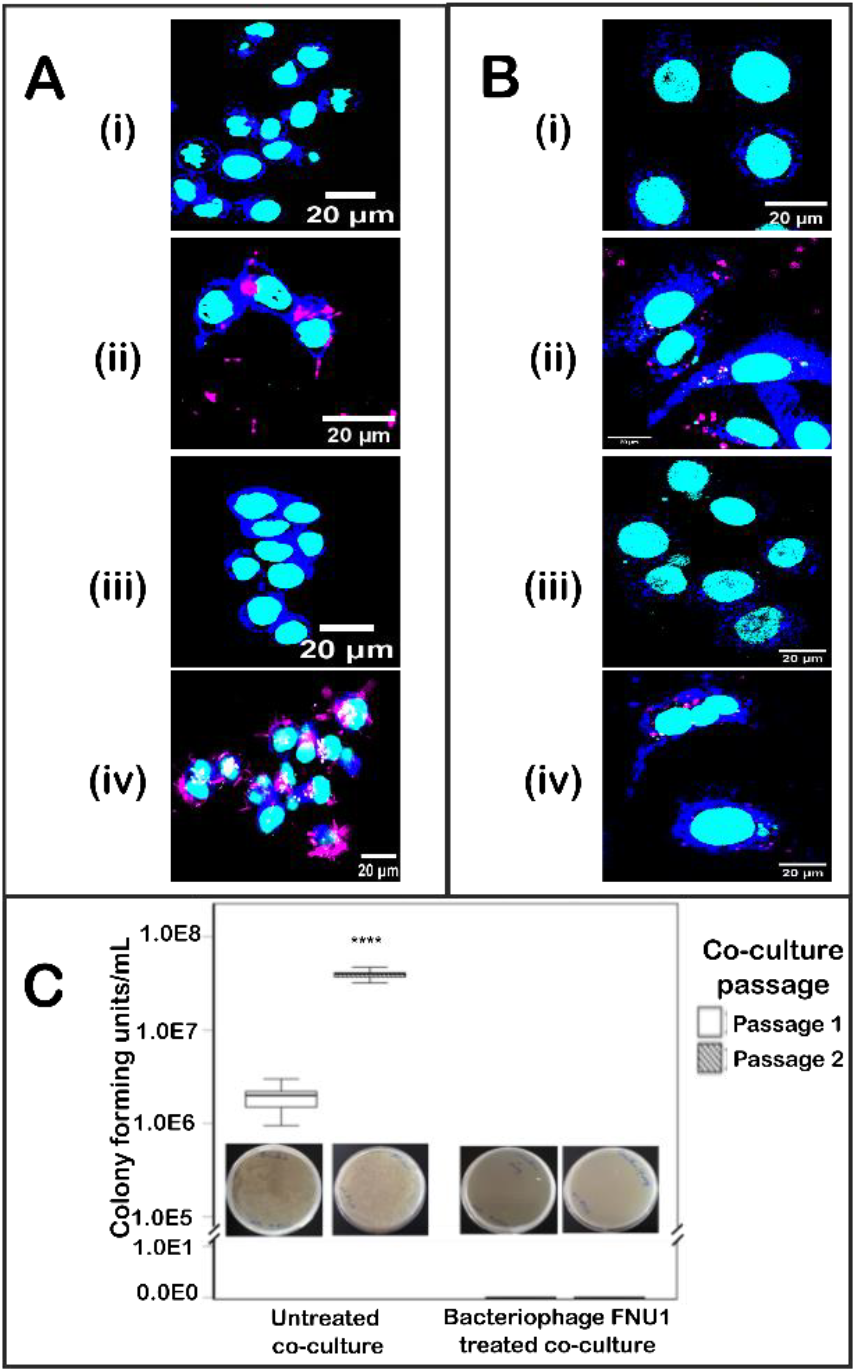
Merged images of the laser scanning confocal microscope showing GI cancer cells HCT116 (A) and MKN1 (B). The GI cancer cells are marked by blue staining of cytoplasm and cyan staining of nucleus (i); GI cancer cells in co-culture with F. nucleatum (magenta) (ii); bacteriophage FNU1 treatment of the co-culture (iii); GI cancer cell/F. nucleatum co-culture treated with 1mg/mL of ampicillin (iv). When the co-culture was passaged, viable bacteria in the untreated co-culture were transferred to a petri dish and showed a significant increase (P < 0.001) in the colony counts. Bacteriophage treated co-cultures did not passage any viable bacteria (C).

### FNU1 bacteriophage treatment reduces *F. nucleatum* driven migration of gastrointestinal cancer cells while antibiotic treatment does not

Infection of GI cancer cell lines HCT116 and MKN1 with *F. nucleatum* significantly increased wound closure compared to the uninfected control group (Figure 3A-D). We next assessed the impact of FNU1 bacteriophage treatment on the migratory potential of infected HCT116 and MKN1 cells. In both cell lines, wound closure was significantly lower at 24, 48 and 72 hours (*p*<0.05, *p*<0.001 and *p*<0.01) compared to infected cells not treated with FNU1 (Figure 3A-D) indicating that FNU1 bacteriophage therapy effectively repressed *F. nucleatum* driven migration of cancer cells. For instance, at 24 hours, wound closure in *F. nucleatum* infected MKN1 cells reached 68 % compared to 45 % in uninfected controls (*p* < 0.05). By 72 hours, infected cells showed complete wound closure (100 %), indicating a significant pro-migratory effect of the bacteria on cancer cells. In contrast, FNU1 bacteriophage significantly prevented these migratory effects. At 24 hours, the wound closure in the FNU1-treated group was 42 %, compared to 68 % in infected cells without FNU1 (*p* < 0.01). At 48 hours, the wound closure was 70 % in the FNU1 treated cells, markedly lower than in the infected group without FNU1 treatment, which had completely closed (*p* < 0.001). There was no significance difference between uninfected cells and infected cells treated with FNU1 bacteriophage. These findings indicated that FNU1 limited the effects of *F. nucleatum*, thereby attenuating the bacterial-dependent increase in cancer cell migration. When a crude preparation of bacteriophage FNU1 was added to the cancer cells, proliferation was significantly increased (*P* < 0.001). However, these proliferative effects were abrogated when the bacteriophages were purified using NaCl/PEG precipitation and Triton^™^ X-100 treatment, removing LPS and other bacterial cellular debris (Figure 3E). There was no significant difference between the proliferation of HCT116 colon cancer cells in monoculture and those co-cultured with *F. nucleatum* treated with purified bacteriophage FNU1 (*p* = 0.072), with purified bacteriophage alone (*p* = 0.272) or with the antibiotic ampicillin (*p* = 0.519). Treatment of the HCT116/*F. nucleatum* co-culture with ampicillin resulted in significantly increased proliferation of the cancer cells (*p* < 0.001) (Figure 3F and G). Apart from the experiments depicted in Figure 3E, all other experiments were performed using only purified FNU1 bacteriophage (rid of LPS and other bacterial cellular debris). The GI cancer cell/*F. nucleatum* cocultures treated with the antibiotic ampicillin did not reduce the migratory effects of *F. nucleatum* (Figure 3 A-D) and resulted in even higher proliferation of the GI cancer cells than those exerted by coculturing *F. nucleatum* alone (Figure 3F and G), *p* < 0.01.

**Figure 3:**
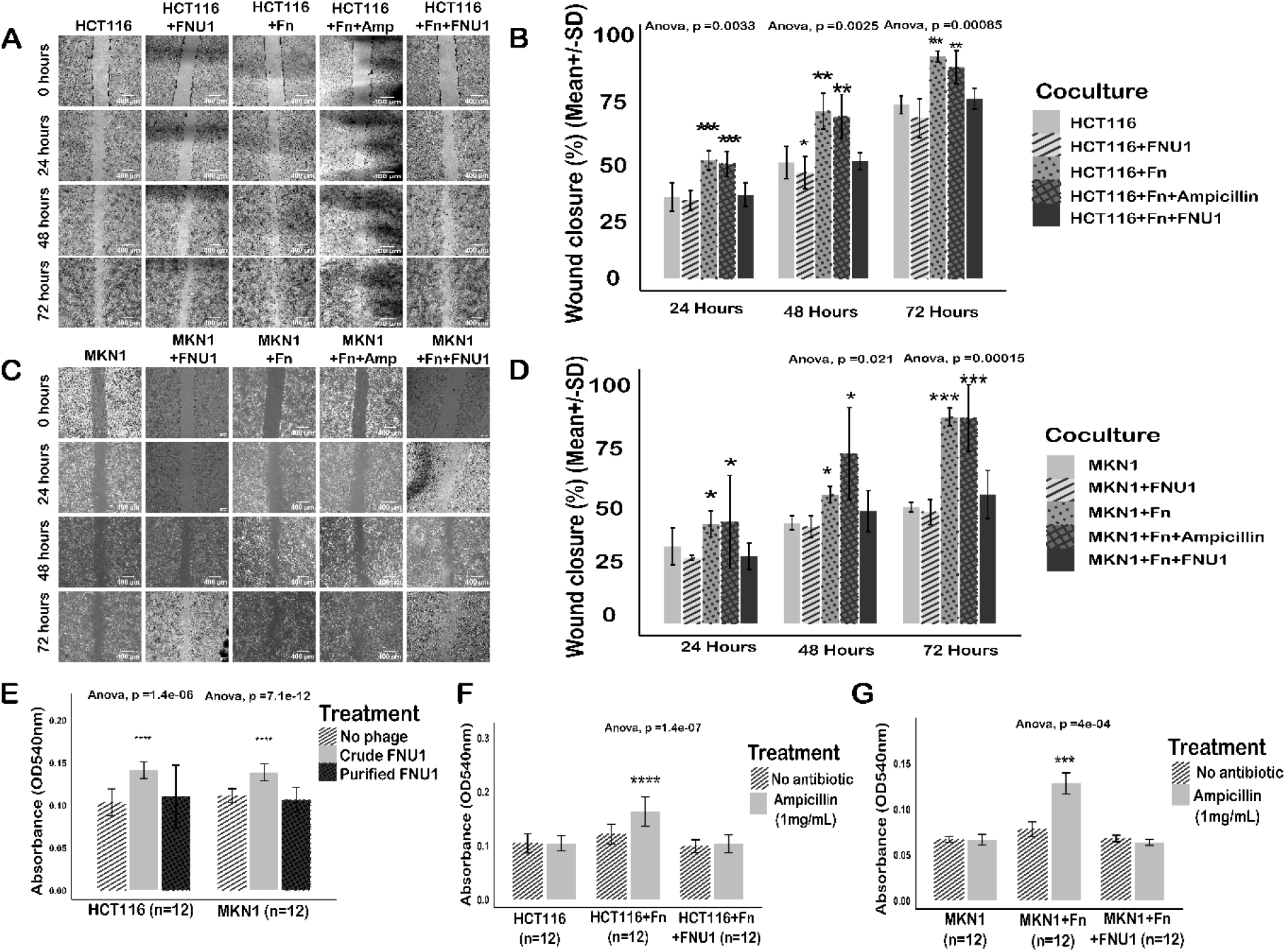
Representative bright field microscope images of wound healing assay for F. nucleatum infected gastrointestinal cancer cells treated with FNU1 bacteriophage. Migration of different co-cultures and treatments for HCT116 (A) and their proportion of wound closure (B). Migration of different co-cultures and treatments for MKN1 cells (C) and their proportion of wound closure (D). * p < 0.05, **p < 0.01 and *** p < 0.001 and ****p < 0.0001; data were analysed by two-way ANOVA, Tukey multiple comparison test, n=3, mean ± SEM, scale bar, 200 µm (E). Proliferation of HCT116 and MKN1 cells treated with crude or purified bacteriophage preparations (F,G). Proliferation of single HCT116 or MKN1 cells or co-cultures with F. nucleatum in the absence or presence of ampicillin; with similar effects to ampicillin treated HCT116/F. nucleatum (F) and MKN1/F.nucleatum (G) cocultures.

### FNU1 negates *F. nucleatum* induced intracellular ROS production in GI cancer cell lines

Intracellular ROS production was measured in HCT116 colon cancer and MKN1 gastric cancer cells in a range of culture conditions, including i) in monocultures ii) co-cultured with *F. nucleatum* iii) co-cultured with *F. nucleatum* and bacteriophage FNU1 iv) with 5-FU and oxaliplatin added to culture conditions i)-iii) above. Over the first 6h of the experiment, there was no statistically significant change in ROS production in HCT116 cells for each category of treatment and culture type (*p* = 0.317; Figure 4A). *F. nucleatum* increased the production of intracellular ROS in HCT116 colon cancer cells (*p* < 0.001) independent of time or chemotherapy treatment. When applied, FNU1 significantly reduced intracellular ROS production in the co-culture (*p* < 0.001), although ROS production was still significantly higher than colon cancer cells on their own (*p* < 0.001). When no bacteria or bacteriophage were added to the colon cancer cells, there was no significant difference in the intracellular ROS production between untreated cells and those treated by 5-FU (*p* = 0.438) or oxaliplatin (*p* = 0.071).

**Figure 4:**
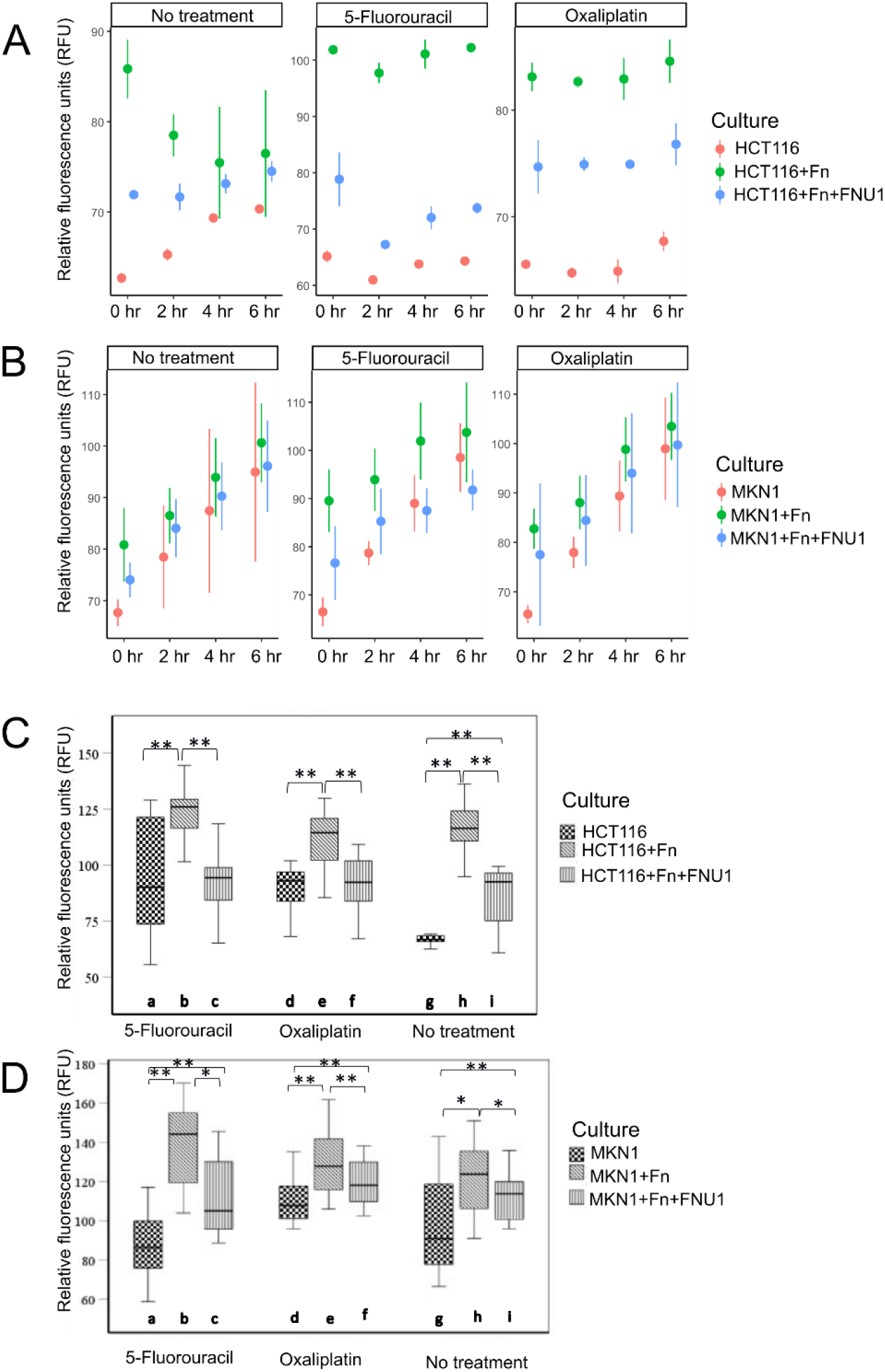
Intracellular ROS production and induction of autophagy after chemotherapy. HCT116 colon cancer cells (A) and MKN1 gastric cancer cells (B) were cultured in the presence of no treatment (left panel), 5-fluorouracil (middle panel) or oxaliplatin (right panel). F. nucleatum increased ROS production in GI cancer cells independent of time or treatment (ANOVA: p < 0.001). Autophagy flux in (C) HCT116 colon cancer and (D) MKN1 gastric cancer cells in monoculture, in co-culture with F. nucleatum, treated with bacteriophage FNU1 and treated with chemotherapy drugs 5-FU and oxaliplatin. Box plots represent the median value and lower 25th and upper 75th quartiles. * Indicates p < 0.05. ** Indicates p < 0.001.

Unlike in colon cancer cells, over the first 6h of the experiment, there was a significant increase in ROS production in MKN1 gastric cancer cells for each category of treatment and culture type (*p* < 0.001, Figure 4B). Similar to colon cancer cells, *F. nucleatum* significantly increased intracellular ROS production in gastric cancer cells (*p* < 0.001) while addition of bacteriophage FNU1 to the co-culture resulted in a significant reduction in intracellular ROS production (*p* < 0.001). Similar to what was seen in the colon cancer cells, when no bacteria or bacteriophage were added to the gastric cancer cells, there was no significant difference in the intracellular ROS production between untreated cells and those treated by 5-FU (*p* = 0.190). There was no significant statistical difference between 5-FU and oxaliplatin treatments (*p* = 0.123).

### FNU1 negates *F. nucleatum* induced autophagy in GI cancer cells

In colon cancer cells (Figure 4C), not exposed to chemotherapy, *F. nucleatum* increased the autophagic flux (*p* < 0.001; Figure 4Ch vs Figure 4Cg) while bacteriophage FNU1 was capable of significantly reducing this effect (*p* < 0.001; Figure 4Ci vs Figure 4Ch), to levels that were still significantly higher than the autophagic flux in untreated colon cancer cells (*p* < 0.001; Figure 4Ci vs Figure 4Cg). *F. nucleatum* increased autophagic flux in both 5-FU (*p* < 0.001; Figure 4Cb vs Figure 4Ca) and oxaliplatin (*p* = 0.001; Figure 4Ce vs Figure 4Cd) treated co-cultures. In these experiments where the drugs were added, FNU1 treatment reduced the autophagic flux to levels similar to the cancer cells without *F. nucleatum* stimulation (5-FU: *p* = 0.727, Figure 4Cc vs Figure 4Ca; oxaliplatin: *p* = 0.267, Figure 4Cf vs Figure 4Cd).

As seen in colon cancer cells, the addition of *F. nucleatum* to cultures of MKN1 cells (Figure 4D) increased autophagic flux (*p* = 0.002; Figure 4Dh vs Figure 4Dg), which was significantly reduced by treating with FNU1 (*p* = 0.005; Figure 4Di vs Figure 4Dh), although these levels were still higher than gastric cancer cells in monoculture (*p* = 0.018; Figure 4Di vs Figure 4Dg). This trend continued when the co-cultures were treated with chemotherapy agents. In MKN1 cell co-cultures with 5-FU or oxaliplatin, autophagic flux induced by *F. nucleatum* (*p* < 0.001; Figure 4Db vs Figure 4Da and Figure 4De vs Figure 4Dd, respectively) was significantly reduced by bacteriophage FNU1 (*p* < 0.001; Figure 4Dc vs Figure 4Db and Figure 4Df vs Figure 4De, respectively) albeit it still remained higher than gastric cancer cells in monoculture (*p* < 0.001; Figure 4Dc vs Figure 4Da and Figure 4Df vs Figure 4Dd, respectively).

### Modulation of apoptosis and impact on GI cancer cell proliferation

Cell death as a result of the co-culture and/or treatment with chemotherapy was calculated as a total proportion of cells stained with annexin V and propidium iodide via flow cytometry. In both colon cancer (Figure 5A and 5B) and gastric cancer cells (Figure 5D and 5E), treatment with either 5-FU or oxaliplatin significantly increased apoptosis regardless of the type of culture (*p* < 0.001). Conversely, co-culture of colon cancer or gastric cancer cells with *F. nucleatum* inhibited apoptosis in the presence of chemotherapy drugs (*p* = 0.004; Figure 5Bb vs Figure 5Ba; Figure 5Be vs Figure 5Bd and Figure 5Eb vs Figure 5Ea; Figure 5Ee vs Figure 5Ed, respectively). This effect that was negated by FNU1 (*p* < 0.001; Figure 5Bc vs Figure 5Bb; Figure 5Bf vs Figure 5Be and Figure 5Ec vs Figure 5Eb; Figure 5Ef vs 5De, respectively).

**Figure 5:**
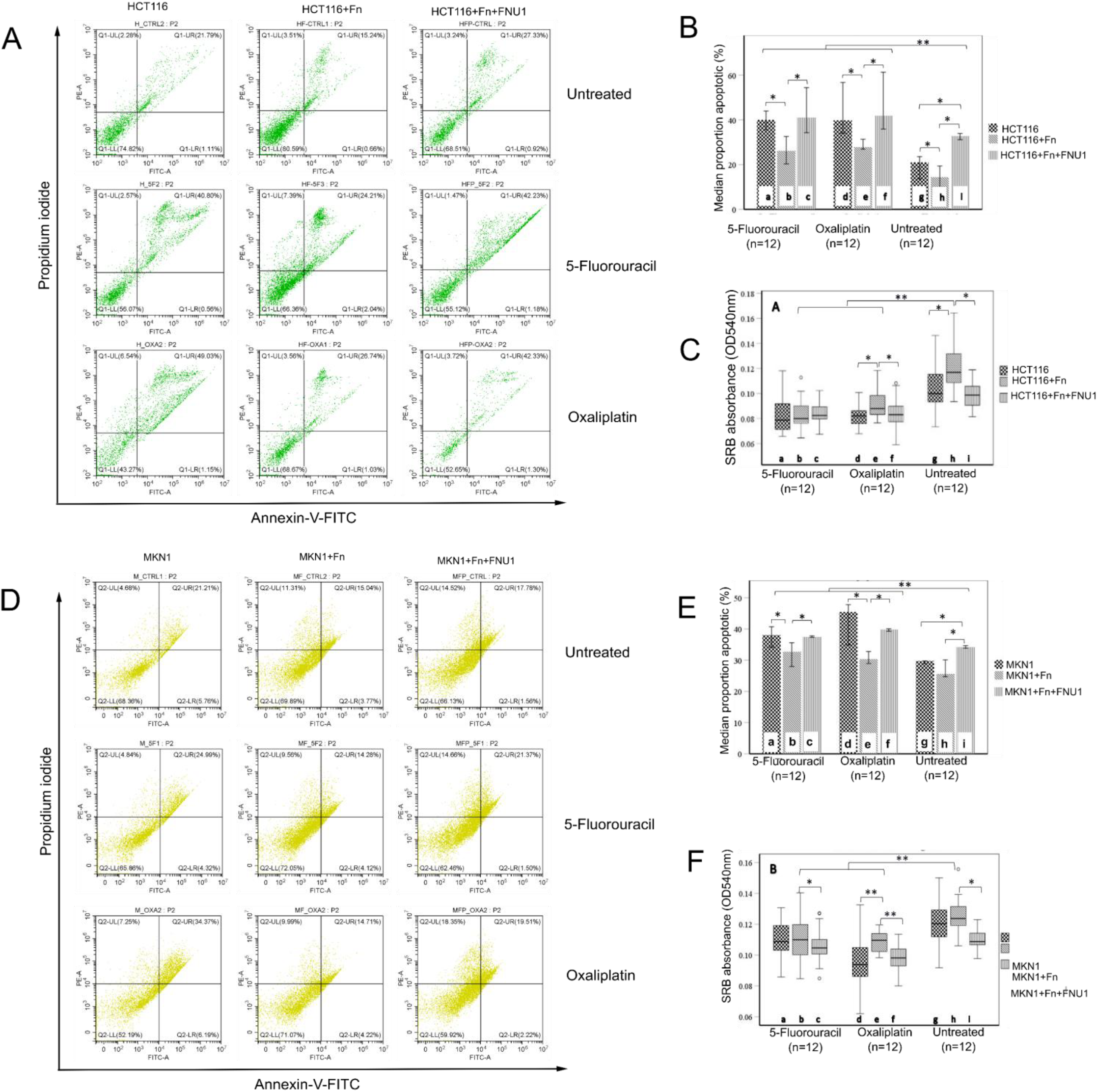
Apoptosis in HCT116 colon cancer cells. A) Flow cytometry scatterplot (B) Flow cytometry data shown as proportion of cells undergoing apoptosis. Apoptosis in MKN1 cancer cells (C). Flow cytometry scatterplot (D) Proportion apoptotic graph. Proliferative/viability effects of F. nucleatum and FNU1 on HCT116 colon cancer (E) and MKN1 gastric cancer cells (F) treated with chemotherapy drugs 5-fluorouracil and oxaliplatin. Box plots represent the median value and lower 25th and upper 75th quartiles. ° indicates an outlier datapoint, * Indicates p < 0.05.** Indicates p < 0.001.

In untreated colon cancer cells, *F. nucleatum* inhibited apoptosis (*p* = 0.046; Figure 5Bh vs Figure 5Bg). In comparison, in untreated gastric cancer cells, the difference in apoptosis between *F. nucleatum*/MKN1 co-cultures and MKN1 cells in monoculture was not significant (*p* = 0.113; Figure 5Eh vs Figure 5Eg). In the absence of drug treatment, addition of FNU1 significantly increased the proportion of cells in apoptosis, when compared to cancer cells co-cultured with *F. nucleatum* (Figure 5 Bi vs 5Bh, *p* = 0.028 and Figure 5 Di vs 5Dh, *p* = 0.026), or in monoculture (Figure 5 Bi vs 5Bg, *p* = 0.028 and 5 Di vs 5Dg, *p* = 0.026) in colon and gastric cancer cell lines, respectively.

Interestingly, bacteriophage FNU1 induced greater levels of apoptosis than 5-FU in co-cultures of *F. nucleatum* and colon cancer cells (*p* = 0.015; Figure 5Bi vs Figure 5Bb) but not in co-cultures of *F. nucleatum* and gastric cancer cells (*p* = 0.394; Figure 5Ei vs Figure 5Eb). Compared to oxaliplatin, bacteriophage FNU1 induced greater levels of apoptosis in both colon and gastric cancer co-cultures (*p* = 0.004; Figure 5Bi vs Figure 5Be and *p* = 0.002; Figure 5Ei vs Figure 5Ee, respectively).

Overall, in colon cancer cells, treatment with 5-FU increased apoptosis from a median of 14.5 % (Figure 5Bh) in untreated *F. nucleatum*/HCT116 co-culture to a median of 26.1 % 5-FU (Figure 5Bb) while in oxaliplatin treated colon cancer cells, apoptosis was increased to a median 27.8% (Figure 5Be). Treatment of HCT116 cells with bacteriophage FNU1 in the absence of chemotherapy increased apoptosis to a median of 32.7 % (Figure 5Bi). The combination of 5-FU and FNU1 in colon cancer cells co-cultured with *F. nucleatum* increased apoptosis to a median of 41.1 % (Figure 5Bc); in similar co-cultures, oxaliplatin with FNU1 increased apoptosis to 41.9 % (Figure 5Bf). Therefore, in HCT116/*F. nucleatum* co-cultures, bacteriophage FNU1 increased apoptosis efficacy of 5-FU from 26.1 % to 41.1 %, while that of oxaliplatin was increased from 27.8 % to 41.9 %.

In gastric cancer cells, the median apoptosis levels were increased from 25.6 % (Figure 5Eh) to 32.7 % (Figure 5Eb) with 5-FU and to 30.4 % with oxaliplatin (Figure 5Ee). FNU1 increased apoptosis to a median of 34.1 % (Figure 5Ei) in MKN1/F. nucleatum co-cultures. The combination of FNU1 with 5-FU in gastric cancer cell co-culture with *F. nucleatum* further increased apoptosis to 37.4 % (Figure 5Ec) and FNU1 with oxaliplatin further increased apoptosis to 39.9 % (Figure 5Ef).

### FNU1 bacteriophage works in synergy with chemotherapeutic agents to negate *F. nucleatum* induced gastrointestinal cancer cell proliferation

Treatment of HCT116 with the chemotherapy drugs (5-FU or oxaliplatin) led to a significant reduction in cell viability, irrespective of whether the cells were co-cultured with *F. nucleatum* or not (Figure 5C). While *F. nucleatum* infection increased proliferation of untreated and oxaliplatin-treated HCT116 cells (5Ee), the addition of FNU1 bacteriophage reverted cell proliferation back to baseline (Figure 5Cf). However, this effect was not observed after treatment with 5-FU, possibly suggesting that 5-FU exerts greater anti-bacterial activity than oxaliplatin under these culture conditions (Figure 5C).

In MKN1 gastric cancer cells co-cultured with *F. nucleatum* in the absence of chemotherapy drugs, there was no significant difference between MKN1 cells in monoculture and those co-cultured with *F. nucleatum* (*p* = 0.253; Figure 5Fg vs Figure 5Fh). However, treatment with bacteriophage FNU1 of *F. nucleatum*/MKN1 co-cultures resulted in significantly decreased proliferation of gastric cancer cells (*p* = 0.002; Figure 5Fi vs Figure 5Fh). With addition of oxaliplatin, *F. nucleatum* significantly increased MKN1 cell proliferation (*p* < 0.001; Figure 5Fe vs Figure 5Fd)) while FNU1 treatment resulted in gastric cancer cell proliferation that was similar to the baseline (*p* = 0.056; Figure 5Ff vs Figure 5Fd). In experiments where 5-FU was added to the culture, *F. nucleatum* did not significantly increase MKN1 cell proliferation compared to those in monoculture (*p* = 0.909; Figure 5Fb vs Figure 5Fa) but treatment of *F. nucleatum*/MKN1 co-cultures with FNU1 resulted in significant reduction in proliferation of the gastric cancer cells (*p* = 0.012; Figure 5Fc vs Figure 5Fa).

## Discussion

The recent recognition of the important role of the microbiome in tumourigenesis has paved the way for its potential manipulation by bacteriophages [9]. In this study, we have used a bacteriophage with selective, lytic activity against *F. nucleatum subsp. polymorphum*, FNU1, to assess efficacy in controlling cancer growth stimulated by this oncobacterium. It has been recognised that *F. nucleatum* binds to Gal-GalNAc through its expression of the Fap2 receptor [37]. Using co-cultures and antibiotic protection assays, we provide data to suggest that *F. nucleatum* uses the Gal-GalNAc receptors not only in *F. nucleatum* attachment, but possibly also for entry into gastric and colon cancer cells, and this protects the bacteria from clearance by ampicillin. *F. nucleatum* increases colon and gastric cancer cell proliferation *in-vitro*, and this oncogenic effect is negated by treatment with FNU1. Our study reveals that *F. nucleatum* promotes spread or invasiveness of cancer cells, increases intracellular ROS production, promotes autophagy and inhibits apoptosis in untreated colon and gastric cancer cells, as well as those treated with chemotherapy drugs, and these effects are also negated/reversed by treatment with FNU1.

The role of *F. nucleatum* in carcinogenesis has been studied, where presence of this bacteria or its DNA is associated with worse outcomes and treatment failures [18,40,41]. The DNA load of *F. nucleatum* in tumour tissues and tumour growth instigated by this bacterium are reduced after treatment with the antibiotic metronidazole in murine models of colorectal [18] and breast cancer [42]. Metronidazole has been shown to be able to penetrate eukaryotic cells and kill intracellular bacteria [43]. However, its use as an adjunct in cancer therapy may be questionable, as it induces DNA damage, as well as dysbiotic effects on the microbiota, and has been associated with cancer progression [44,45]. Similar to our findings, a recent study has used gentamicin to show that *F. nucleatum* enters cancer cells [46]. Ampicillin does not enter eukaryotic cells [47], and in our study, not only failed to clear intracellular bacteria but promoted the overgrowth of the cancer cells in co-culture with *F. nucleatum*. We have shown that bacteriophage FNU1 is capable of eliminating intracellular *F. nucleatum*. Previously, bacteriophages have been shown to eliminate intracellular bacteria when internalised through endocytosis by phagocytic cells [48,49] or carried into the eukaryotic cell by bacteriophage infected obligate intracellular bacterial cells [50]. Bacteriophages on their own have also been shown to interact with and traverse non-phagocytic epithelial cells through their receptors [51].

*F. nucleatum* has been previously reported to increase cancer cell proliferation *in vitro* by increasing ROS production [15], induction of autophagy [13] and inhibition of apoptosis [12,52]. Other oncobacteria such as *Enterococcus faecalis* have also been shown to increase ROS production in co-culture with mammalian cells [34,53-55]. Bacteriophage EFA1 treatment of *E. faecalis* in co-culture with colon cancer cells *in vitro* resulted in a further increase of ROS production [34]. In contrast, our findings here show that FNU1 significantly reduced the ROS produced when colon cancer and gastric cancer cells were grown with *F. nucleatum*. Other studies involving immune cells [56-58] showed a reduced ROS production upon bacteriophage treatment, similar to findings from this study. Despite the varying responses to ROS production, both FNU1 and EFA1 bacteriophage treatments resulted in overall decrease in proliferation of cancer cells. Further elucidation of the exact role of ROS in these culture systems is required.

As shown in our study, and by others [12,13], *F. nucleatum* reduces the sensitivity of cancer cells to chemotherapy drugs such as 5-FU and oxaliplatin, when measured as a proportion of apoptotic cells. We could not find other studies that employed SRB assays to define cancer proliferative effects of *F. nucleatum* after chemotherapy drug treatments. Studies exploring the proliferative effects of 5-FU on *F. nucleatum* infected cancer cells have used CCK8 assays [12,13] or MTT and MTS assays [59], which are commonly used as a measure of cell cytotoxicity [35], and like apoptosis assays, are dependent on and measure the quantity of viable cells [60]. SRB assays measure cell mass regardless of viability [35,61].

To investigate whether *F. nucleatum* reduced cytotoxicity of 5-FU and oxaliplatin, and if bacteriophage FNU1 was able to negate such effects, we investigated induction of autophagy. Autophagy is a well characterised mechanism by which *F. nucleatum* induces chemotherapy resistance [13,15]. We were able to demonstrate that FNU1 was able to significantly reduce the autophagic flux resulting from *F. nucleatum* stimulation of cancer cells in co-culture. We could not find any other study that has investigated bacteriophage capacity to modulate autophagic flux.

Finally, we showed that the chemotherapy resistance induced *by F. nucleatum* through inhibition of apoptosis may be negated by bacteriophage treatment. Previously, bacteriophages against *F. nucleatum* have been used to augment cancer therapy. Although apoptosis was not measured, Zheng and colleagues performed transcriptomic studies showing that elimination of *F. nucleatum* using bacteriophages inhibited the upregulation of anti-apoptotic genes and downregulated pro-autophagy genes [62]. In our study we quantified this effect and demonstrated a synergistic activity of FNU1 with chemotherapy agents.

We demonstrated an increased proliferative effect of *F. nucleatum* in co-cultures with HCT116 colon cancer and MKN1 gastric cancer cells and when these co-cultures were treated with oxaliplatin but not 5-FU. 5-FU is a potent antimicrobial compound through its inhibition of DNA synthesis and induction of DNA damage in bacterial cells [63]. Oxaliplatin on the other hand does not appear to have significant antimicrobial activities in cell culture systems [64]. However, it is important to note that these are purely experimental models, and in a complex tumour microbiome comprising many other bacterial species, the structure and function of cancer drugs such as 5-FU can be altered. For instance, pks+ *Escherichia coli* can modify the chemical structure of 5-FU through converting uracil into 5,6-dihydrouracilin de-activating 5 - FU in the process [65].

## Conclusion

Our study showed that using bacteriophage FNU1 to control *F. nucleatum* limited the proliferation and migration of cancer cells, increased their chemosensitivity, and reduced the inflammatory burden by limiting intracellular ROS production. Further, FNU1 was capable of reducing autophagic flux induced by *F. nucleatum* which is key to preventing chemotherapy resistance [13]. These effects led to an increase in the efficacy of chemotherapy in both colon cancer and gastric cancer cells co-cultured with *F. nucleatum*, as measured by the proportion of apoptotic cells. The results suggest that further testing of bacteriophages against oncobacteria using *in vivo* models is warranted in order to manipulate the microbiome and enhance chemotherapeutic efficacy against tumours colonised by these cancer promoting microbes.

## Acknowledgements

We acknowledge the Australian Cancer Research Foundation and The Collie Foundation for providing funds to purchase the Zeiss 980 confocal microscope at the Olivia Newton-John Cancer Research Institute

## Author roles

Conceptualization: JT, MK

Investigation: MK, SS, BJ

Methodology: MB, JT, MK

Visualization: MK, SS, BJ

Supervision: JT, MB

Writing - original draft: MK, BJ

Writing – review & editing: MB, JT, MK

## Data availability

Samples of Fusobacterium bacteriophage FNU1 can only be shared by the authors for non-commercial purposes and under material transfer agreement. Readers wishing to obtain them should contact Latrobe University Commercialisation Office.

